# RTM-align: an improved RNA alignment tool with enhanced short sequence performance via post-standardization and fragment alignment

**DOI:** 10.1101/2024.05.27.595311

**Authors:** Zijie Qiu, Sheng Xu, Junkang Wei, Tao Shen, Siqi Sun

## Abstract

Understanding the three-dimensional structure of RNA is crucial for studying various biological processes. Accurate alignment and comparison of RNA structures are essential for illustrating RNA functionality and evolution. The existing RNA alignment tools suffer from limitations such as size-dependency of scoring functions and inadequate handling of short RNA fragments, leading to the conflicting interpretation of structural and functional relationships among RNA molecules. Hence, we introduce RTM-align, a novel RNA structural alignment tool enhanced for short RNAs. RTM-align employs the RTM-score, which integrates post-standardization to ensure size-independence and utilizes a fragment alignment strategy that improves alignment accuracy by concentrating on structural motifs and local structural elements. Benchmarking results demonstrate that RTM-align outperforms existing tools in RNA structure comparison, offering a universal scoring scale regardless of RNA length. The improvement is particularly evident in evaluating predicted structures for CASP15 RNA targets, with significant enhancements observed for the short RNA target R1117. RTM-align is expected to significantly improve the accuracy and reliability of RNA structure alignment and comparison, thereby aiding in the deeper understanding and discovery of novel RNA functions and their interactions in biological systems. RTM-align is now available at https://github.com/BEAM-Labs/RTM-align.

## Introduction

RNAs are essential in various cellular activities, acting not only as genetic messengers but also as regulators and catalysts of biochemical processes (1–4). The sequence alignment of biological molecules such as proteins, DNA, and RNA is crucial for elucidating their structure and function (5). For example, AlphaFold (6, 7) employed the massive multiple sequence alignment (MSA) for protein structure prediction. RNA-MSM (8) utilized the homologous sequences from RNAcamp to capture the evolutionary information. Unlike proteins, RNA sequences are generally less conserved (9). However, recent studies have shown that certain RNAs with similar functions, such as tRNA and rRNA, exhibit conservation in structure despite having distinct sequences (10). This underscores the importance of structural alignment in elucidating RNA function (11, 12). The functionality of RNA is intricately linked to its complex three-dimensional structure. Accurate structural alignment of RNA is crucial because structurally similar RNAs often exhibit similar functions, enabling researchers to infer functional relationships and evolutionary conservation (13).

Given the critical link between structure and function of RNA, a bunch of RNA-centric structure alignment programs have been developed to compare different RNA structures. These include ARTS (14), STAR3D (15), SARA (16), and Rclick (17). Specifically, some RNA alignment programs utilized TM-score (18, 19) and adapted it for RNA structure alignment. These programs have demonstrated effective performance primarily because of TM-score’s advantageous property of size independence. This feature facilitates meaningful and flexible comparisons of similarity between structures of different lengths. Among these, RMalign (20) and RNA-align (21) are notable representatives. However, both methods exhibit certain limitations, particularly with short RNA sequences, which play a critical role in various diseases(22). RMalign introduced a learnable bias *h* to normalize scoring scales across diverse sequence lengths. However, this method shows limited effectiveness on short sequences and potentially yields higher scores towards them due to deviations from the standard scaling parameter *d*_0_. This undermines the TM-score’s inherent advantage of size independence. In contrast, RNA-align asserted a general form of the function *d*_0_(*L*) and optimized for scale consistency across varying lengths. Though RNA-align achieved a uniform scale in TM-score, it exhibits significant limitations, including unfairness and extreme sensitivity on short sequences. These issues lead to poor performance when assessing homology in short RNA pairs and a low correlation with other structural similarity assessment metrics, such as the Local Distance Difference Test (lDDT) (23), particularly for short RNA targets. This is well illustrated by the observation that some RNA pairs have significant structure coverage but with low TM-scores (<0.5). For instance, 1Y3S_B and 2FD0_B have a TM-score of only 0.3944, despite an RMSD of merely 1.09 Å. Similarly, 6GTD_B and 6GTC_B have a TM-score of 0.3490, while the RMSD is 1.53 Å(Fig. 1).

**Fig. 1.**
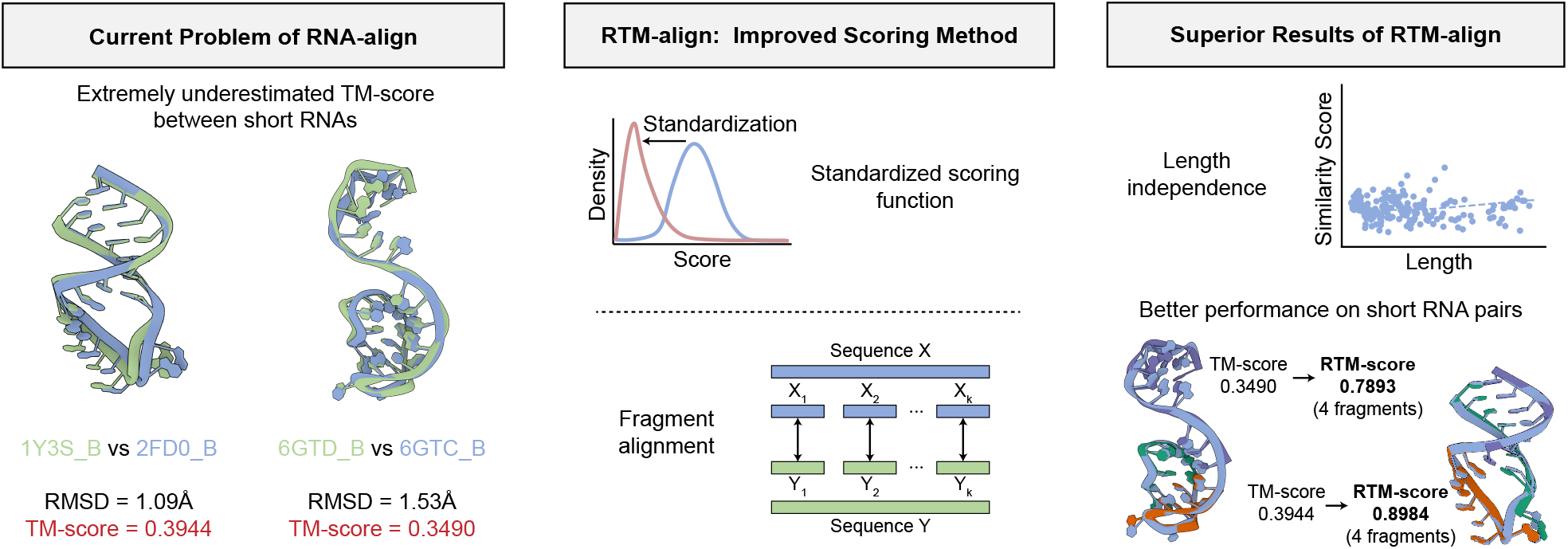
Figure abstract of our work.

In response to these challenges, we developed RTM-align, a novel RNA alignment tool specifically designed to improve the accuracy and reliability of short RNA structure alignments and maintain the capability to handle RNAs of various lengths. RTM-align employs the RTM-score, which utilizes a post-standardized scoring function that is independent of RNA length. This feature enables fairer structural comparisons by eliminating the bias introduced by the varying lengths of RNA molecules. Additionally, RTM-align employs a fragment alignment strategy, focusing on structural motifs and critical local structural elements, which is particularly effective for short RNAs. This approach enhances the alignment accuracy by preventing significant score fluctuations caused by several tiny local deviations. Benchmarking results suggest that RTM-align outperforms existing tools in alignment precision and template scoring, particularly in a CASP15 RNA target. This performance enhancement is critical for applications involving the rapid and precise alignment of short RNAs, which is often required in large-scale RNA structural studies and the exploration of RNA’s role in complex biological systems. Through the capabilities of RTM-align, we aim to provide a robust tool and advance the field of RNA structural analysis.

## Materials and methods

### Fitting the template modeling function for RNA

RTM-align is predicated on the TM-score (18), which was originally developed to assess the quality of protein structure templates and later adapted for the structural alignment of biomolecules such as protein(19) and RNA(20, 21). The TM-score has several advantageous properties as a measure of structural similarity, including its size-independent scale and its ability to compare structures of different lengths. TM-score conforms to the following formula:

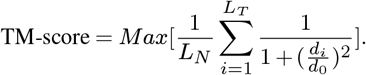

Here, *L*_*N*_ represents the length of the target structure, *L*_*T*_ denotes the length of aligned residues, *d*_*i*_ is the *L*_2_ distance between the *i*-th aligned residues, and *d*_0_ is a scaling factor designed to mitigate the statistical differences in the calculated scores across various lengths. Therefore, the key component of TM-score function is the fitting of the reliance of *d*_0_ on length. According to TM-align (18), *d*_0_ is assumed to be proportional to the average Root Mean Square Deviation (RMSD) for random pairs of structures of similar length, as expressed below:

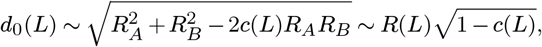

where *R*(*L*) indicates the expected radius of gyration for a structure of this length and *c*(*L*) represents the average correlation coefficient (24) for randomly associated structure pairs of the same length.

Following this methodology, we initially determined the functions of *c*(*L*) and *R*(*L*) using the random pairs dataset (Supplementary Fig. S1). The function *c*(*L*) was fitted on the 6,343 unique RNA structures contained in the dataset, and function *R*(*L*) was derived from the 13,308,344 RNA pairs whose sequence identity falls below 0.4. To align with previous studies, we set the coefficient in *d*_0_(*L*) to 1, establishing the following relationship:

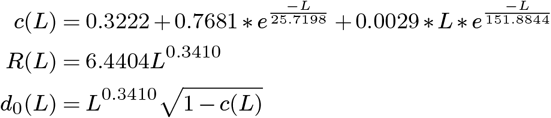

Our RNA alignment program, RTM-align, is based on the *d*_0_ function. In light of the methodological requirements for constructing structure alignments, we adapted the aligning program and parameter settings from RNA-align (21) as our basic alignment method. The alignment procedure involves generating initial alignments between two structures using gapless sliding. This is followed by dynamic programming on the secondary structure, among other techniques, and concludes with iterative refinement based on the computed TM-score. On the other hand, we additionally introduce a fragment alignment procedure for short RNAs, which will be discussed later.

### Post-standardization

RNA-align’s poor performance on short RNAs could partly be attributed to the potential bias toward shorter RNAs when fitting the function of *d*_0_ under the objective of scale consistency across different lengths. In contrast, the traditional method used in RMalign fails to reach consistency partly because there is only one hyperparameter to alleviate scale dependence in the TM-score function. This approach is insufficient for managing the sensitivity associated with the small value of *d*_0_ at short lengths. Therefore, instead of eliminating TM-score’s dependency on length by introducing bias or optimizing on other hyperparameters as TM-align (19), RNA-align (21) or RMalign (20), we prioritize the absolute fairness during TM-score calculation. To achieve this, we introduce a post-standardization procedure to normalize the TM-score. This procedure is conceptually similar to that used in SP-align (25), but employs z-score as the standardization method.

We applied our alignment program with the derived *d*_0_(*L*) on all RNA pairs in the random pairs dataset and calculated the mean and standard deviation of the resulting TM-scores for different lengths. We found that both the mean and standard deviation curves exhibit an inverse proportionality trend (Supplementary Fig. S2). By fitting with the general form 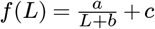, we get:

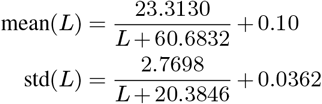

The TM-score of aligned structures on each length, calculated by our program, can be standardized to a Gaussian distribution, which is the classical procedure of z-score. In order to ensure the output TM-score falls within 0 and 1, we apply a sigmoid function to the standardized score, as the following formula indicates:

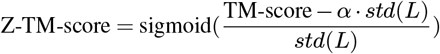

We call the resulting score as RTM-score_BA_ and the aligning program using it as RTM-align (w/o fragment alignment). Note that we first standardize TM-score to a Gaussian *N* (− *α*, 1). The positional hyperparameter *α* is introduced because the mean and standard deviation statistics were calculated on random pairs, which typically have low similarity scores. Additionally, the sigmoid projection is nonlinear. By setting the position center to a negative value, RTM-score_BA_ for more similar pairs will exhibit a near-linear relationship with length. In our experiments, we found that *α* = 2 is an appropriate setting (Supplementary Fig. S4).

### Fragment alignment

Some researchers have introduced fragment alignment (26) to construct initial structure alignments, but the final TM-score is calculated by treating the aligned pairs of two structures as two separate wholes. According to our observations, some short RNA pairs between 20 and 50 nucleotides may show highly similar sequences, but yield exceptionally low TM-scores, as shown in the hard-pairs dataset.

One explanation might be that although two RNA sequences differ only at a few individual nucleotides, these specific variations result in a slight misalignment in their overall three-dimensional structures, leading to a relatively high RMSD. When the sequence length is short (20-50 nt), such deviations can cause a significant reduction in the calculated TM-score. This is because the *d*_0_ value is very small at this length, thus the TM-score is highly sensitive to changes in *d*_*i*_. However, to human experts, these sequences are considered highly similar and can be used as each other’s templates. Therefore, alignment programs should be equipped to avoid situations where minor base discrepancies lead to major global effects, especially when dealing with short sequences.

To address this issue, we propose the incorporation of fragment alignment as a numerical adjustment tailored for short RNAs which takes local alignment performance into consideration. However, it’s crucial to emphasize that the ultimate output of structure alignment is still basic alignment despite the utilization of fragment alignment. The implementation of fragment alignment proceeds as follows:

**Step 1** Given two structures *X* and *Y*, with lengths *L*_*X*_ and *L*_*Y*_ respectively, first pass the two structures into RTM-align (without fragment alignment) to obtain the basic alignment. Extract the aligned structures *X*^*′*^ and *Y* ^*′*^, both with length *L*_*T*_.

**Step 2** Assign a fragment length *L*_*frag*_ (4 or 8). Split *X*^*′*^ into fragments *X*_1_, *X*_2_, …, *X*_*k*_, where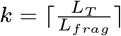. Perform the same splitting for *Y* ^*′*^.

**Step 3** Align each fragment pair *X*_*i*_ and *Y*_*i*_ using the Kabsch algorithm.

**Step 4** Calculate the alignment score for each fragment pair and sum these scores with appropriate weights.

**Step 5** Apply the post-standardization procedure to the score obtained in Step 4 to get the final RTM-score_FA_. Step 4 and step 5 can be more clearly described with the following formula:

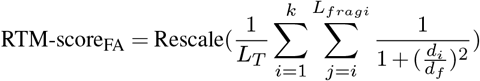

Here, *d*_*f*_ is a normalizing factor for fragment alignment. Setting *d*_*f*_ directly to *d*_0_ for the given length is problematic because aligning two three-dimensional structures is not equivalent to aligning their fragments, which is relatively easier. Additionally, from TM-align, we know that the expected RMSD between short fragments is typically lower than that of complete structures. These factors can lead to an inflated calculated score. This contradicts our intention of using fragment alignment only as a remedy in extreme cases with a few significant misalignments. Therefore, we set *d*_*f*_ to *βd*_0_, where *β* is a hyperparameter. The purpose of setting *β* is to penalize fragment alignment, ensuring that it does not produce a higher score than the original alignment for typical RNA pairs. We have chosen the Rfam-50 dataset as a benchmark to determine *β* because it includes a diverse mix of similar and dissimilar pairs.

*β* should be as high as possible as long as it satisfies the following two constraints:

1. The average RTM-score_FA_ should not exceed the RTM-score_BA_ obtained by using post standardization alone.
2. No more than 5% of the pairs belonging to different families should have a RTM-score_FA_ higher than RTM-score_BA_.

We set the hyperparameter *L*_*frag*_ = 8, and our experiments show that *β* = 0.6 is the best under this condition. As shown in Supplementary Fig. S3, when *β* = 0.6, the average of RTM-score_FA_ is lower than RTM-score_BA_ at each length. While 4.7% of negative pairs in Rfam-50 have higher RTM-score_FA_ than RTM-score_BA_. These are then chosen as our default settings. An alternative is to set *d*_*f*_ to a fixed value and optimize as above.

### RTM-align program

The final RTM-align algorithm incorporates both post-standardization and fragment alignment techniques. Initially, the algorithm calculates a basic TM-score for RNA structures using the fitted *d*_0_. Post-standardization is then applied to rescale the score. Fragment alignment is specifically employed for RNA structures whose lengths range from 20 to 50 nucleotides. In these cases, the greater value between RTM-score_BA_ and RTM-score_FA_ is selected as the final score (Fig. 3a). It is important to note that fragment alignment serves merely as a numerical adjustment to provide fairer scores for short RNAs, rather than a genuine ’alignment’. The final output alignment of the input structure pair will still result from the basic alignment.

### Random pairs dataset

To accurately fit the scaling factor *d*_0_, we built a dataset of RNA pairs with low sequence similarity, as the previous TM-score-based methods did (21). We started by extracting all RNA chains from the Protein Data Bank (PDB) up to May 3rd, 2023, total 18,803 sequences. From this pool, we removed sequences with over 10% unmodeled nucleotides and retained only one PDB file per RNA sequence in cases of multiple PDB structures corresponding to the same sequence. Subsequently, we excluded structures whose PDB contents were inconsistent with the labeled RNA sequences or whose lengths were above 1000 nucleotides. These procedures further reduced the pool to 6,343 RNAs. Although this refined dataset could generate around 20 million pair-wise combinations, we specifically selected pairs with less than 40% sequence identity to focus on dissimilar pairs for better *d*_0_ estimation. This selection process resulted in a final dataset comprising 13,308,344 RNA pairs.

### Rfam-50/Rfam-1000 dataset

The classification threshold for determining whether an RNA structure pair is similar is based on the distribution of TM-scores within a certain dataset. The pairs in this dataset should have labels indicating whether the pair is similar or not. Then, the posterior probability of whether one RNA pair is similar based on its TM-score can be calculated, thus determining the threshold of judging an RNA pair as similar. RNA-align uses the Rfam dataset (27) as the fundamental dataset, while RMalign uses the FSCOR dataset (28, 29). The former clusters RNAs based on sequence identity and structure similarity, while the latter uses RNA functions.

Our work used the Rfam dataset following RNA-align(21). Our data of the mapping between Rfam and PDB were extracted from (30), and we only retained PDB structures that were uniquely assigned to a single Rfam family and had an unmodeled ratio below 0.1. All RNAs selected were paired in an all-to-all manner. A pair is labeled as positive if the two RNAs in it belong to the same Rfam family and vice versa.

Moreover, our work emphasized solving previous methods’ problems on short sequences. Thus we constructed an Rfam dataset that only contains structures with lengths ranging from 20 to 50 nucleotides and excluded families with fewer than 6 structures. Ultimately, this dataset encompassed 308 distinct RNA structures across 21 Rfam families. We named this dataset Rfam-50.

In contrast, a dataset containing both the short and the long RNAs, named Rfam-1000, was constructed as well to make a fair comparison. Rfam-1000 was specifically curated by excluding structures with lengths under 20 or over 1000 nucleotides. Furthermore, families comprising fewer than 10 structures were removed. As a result, Rfam-1000 comprised 1338 distinct RNA structures distributed across 26 Rfam families.

### Hard-pairs dataset

To demonstrate the performance of RTM-align in aligning structurally similar short RNAs misclassified by RNA-align, we built a dataset of pairs having low TM-score but high sequence and structural similarity. We obtained all available RNA structures from the PDB, ranging in length from 20 to 50 nucleotides. All pairwise combinations of these RNAs were then aligned using RNA-align. To identify previously misclassified pairs, we selected the pairs that fulfill these conditions:

1. TM-score below 0.5.
2. Alignment ratio of nucleotide (including pairs within a distance of less than 5.0 Å and other aligned nucleotides) over 90%.
3. Aligned RMSD less than 2 Å.
4. Sequence identity over 80%.

The resulting dataset included 2,882 RNA pairs.

### R1117 dataset

The Critical Assessment of Structure Prediction (CASP) serves as a global benchmark for evaluating protein structure modeling methods. 12 RNA structures were evaluated during CASP15, including a short PreQ1 class I type III riboswitch, designated as R1117, comprising 30 nucleotides. Das et al. (31) highlighted discrepancies in the assessment metrics for R1117, noting that the TM-score values were significantly lower compared with the canonical alignment metrics, including Global Distance Test - Total Score (GDT_TS) (32) values and had low correlation with other metrics such as global Root Mean Square Deviation (RMSD) and local Distance Difference Test (lDDT).

To investigate potential enhancements of RTM-score in its role as a tool for evaluating predicted RNA structures, we established the R1117 dataset. This dataset comprises 153 predicted structures of R1117 submitted during the CASP15 competition, each paired with the reference R1117 structure and evaluated using official metrics including GDT_TS, RMSD, lDDT, and TM-score.

## Results

### RTM-score is a size-independent metric to evaluate RNA structure similarity

In order to examine RTM-score’s scale consistency across the length, we obtained the average output score of RNA-align, RMalign and RTM-align on each given length respectively. These values were derived from analyzing 16,280,069 RNA pairs within the random pairs dataset. When aligning a shorter RNA with a longer one, the score corresponding to the shorter RNA is selected, and the pair is categorized under the shorter length. Fig. 2a presents the average score as a function of RNA lengths for RNA-align, RMalign, and RTM-align. The scores generated by RNA-align and RTM-align are more independent of length, in contrast to those from RMalign, which demonstrate a significant declining trend within the short interval of 20-200 nucleotides. In order to quantify the relationship between score and length, particularly for short sequences, we compared the absolute values of the slopes of the TM-score and length across a specified range (Supplementary Table.S1). A lower value suggests greater size-independence. It is observed that RMalign displays higher values than RNA-align and RTM-score across each length interval between 20 to 200 nucleotides.

**Fig. 2.**
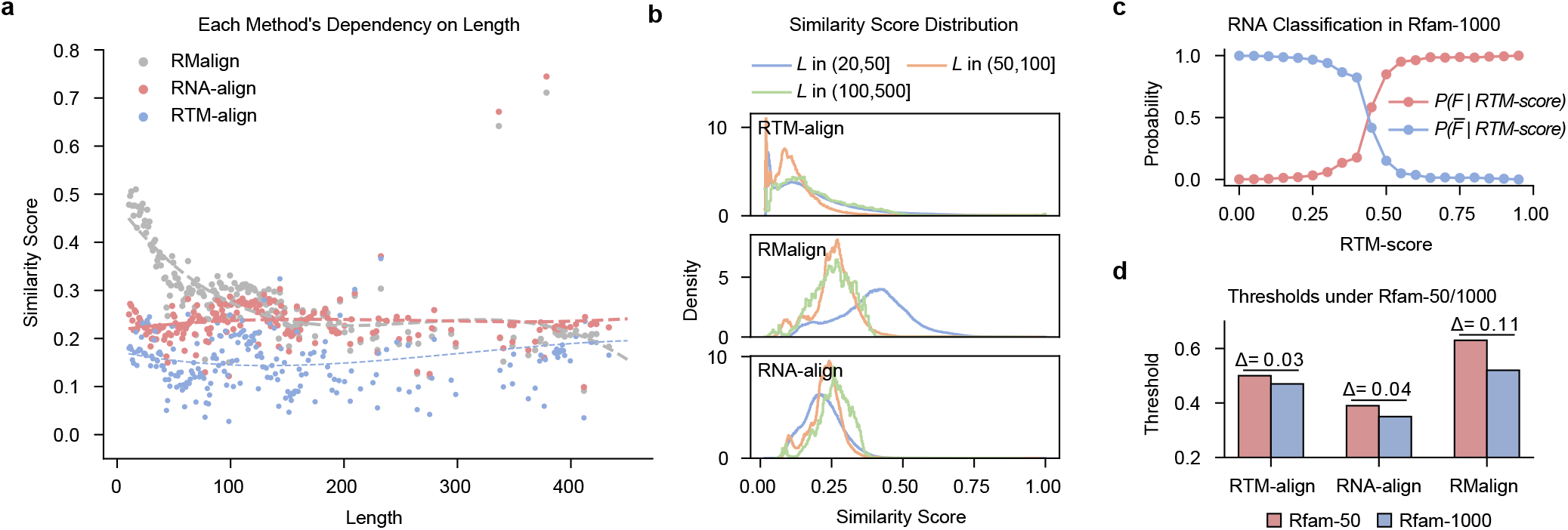
**(a)** Dependency of each method on sequence length. Each data point represents the average similarity score for a given sequence length. **(b)** Distribution of similarity scores among RNA pairs within the random pairs dataset, categorized by length intervals of (20, 50], (50, 100], and (100, 500], respectively. **(c)** Posterior probability of an RNA pair being classified into the same or different Rfam families based on the RTM-scoreBA. **(d)** Each method’s classification threshold under Rfam-50 and Rfam-1000.

**Fig. 3.**
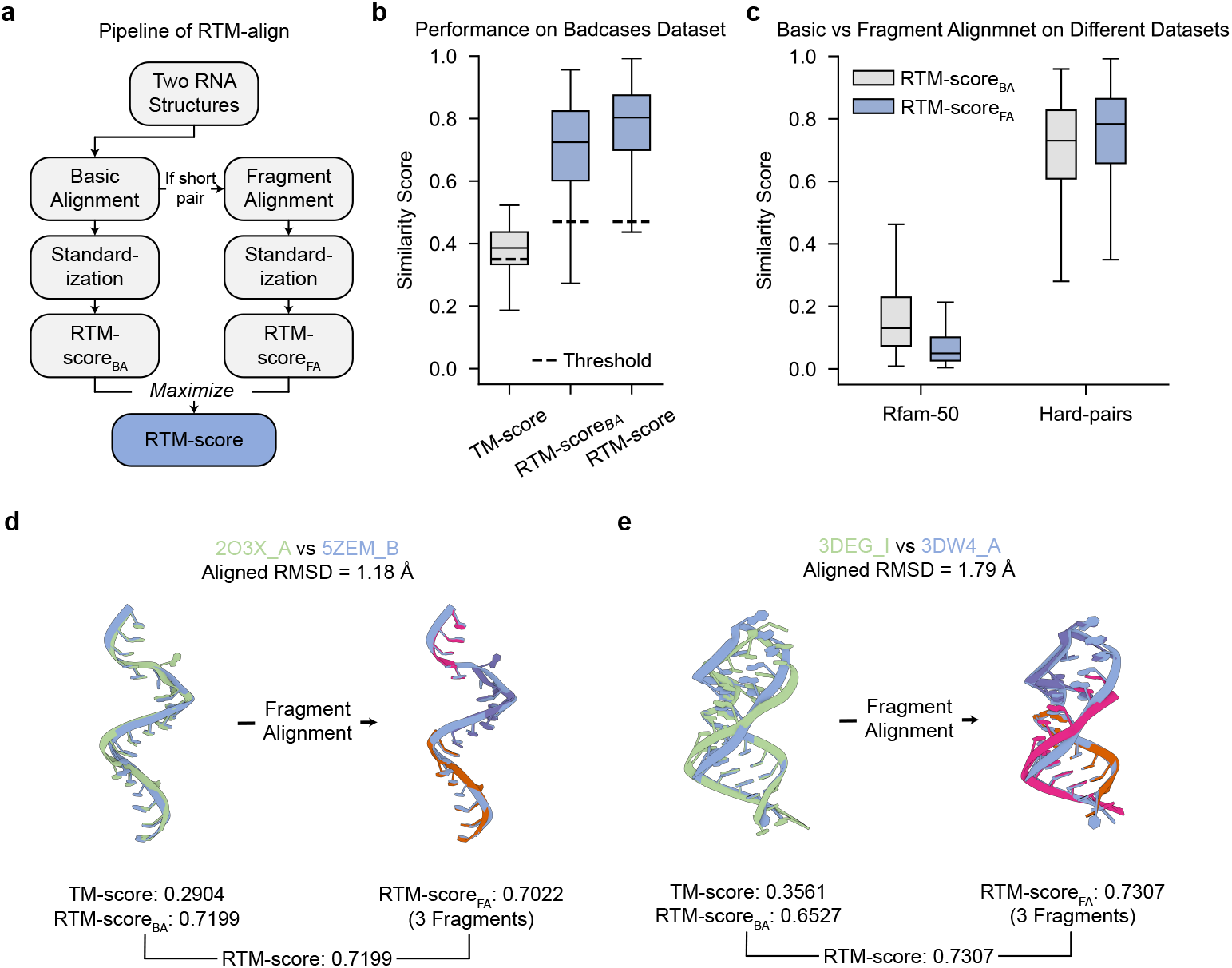
**(a)** Pipeline of RTM-align **(b)** Distribution of TM-score and RTM-score on the hard-pairs dataset. **(c)** Comparison of RTM-align using basic alignment or fragment alignment on the Rfam-50 dataset and the hard-pairs dataset. **(d)** TM-score and RTM-score of RNA pair 2ET3_A and 4GPW_A. **(e)** TM-score and RTM-score of RNA pair 3DEG_I and 3DW4_A.

Given that RMalign tends to yield higher scores for short RNAs, it loses the inherent advantage in the TM-score, which is capable of providing a uniformly scaled and equitable score across structures of varying lengths. This limitation restricts the potential application of RMalign as a standard structural similarity scoring tool. To elucidate this, we analyzed the distribution of TM-score across various RNA length intervals, specifically (20, 50], (50, 100], and (100, 500]. Fig. 2b illustrates a notable shift in the RMalign output towards higher values for the shortest length range (20, 50], as compared to longer intervals, while RNA-align and RTM-align maintain a consistent output distribution across all three length ranges.

Similar to RNA-align (21), we evaluated the reliability of the program by examining its performance in the Rfam family assignment. We statistically analyzed the posterior probability of an RNA pair belonging to the same family based on its TM-score using the Rfam-50/Rfam-1000 dataset. This allowed us to compare the programs’ performance on short sequences and on all sequences separately. Initially, RNAs in the dataset were compared in an all-to-all manner. Based on the computed TM-score and their family affiliations, we plotted the posterior probability of a pair being in the same family against given TM-score values (Fig. 2c). This curve exhibits a sigmoid-like trend. The intersection point of the two curves was designated as the classification threshold. To calculate classification accuracy, pairs with a TM-score below the threshold were labeled as belonging to different families, and vice versa. As indicated in Table.1, RNA-align’s classification accuracy on Rfam-50 and Rfam-1000 are 0.9284 and 0.9911 respectively, which are both surpassed by RTM-score’s 0.9363 and 0.9927, while using fragment alignment or not only has a slight impact on this task. This suggests that RTM-align is more accurate than RNA-align in the Rfam family assignment task.

Furthermore, RMalign exhibits the largest discrepancy in the Rfam classification threshold (0.11) when computed for Rfam-50 versus Rfam-1000, in contrast to the smaller, consistent difference of 0.04 observed with RNA-align and RTM-align (Fig. 2d). These findings highlight the negative impact of RMalign’s preference for short sequences on fairness in family classification tasks. On the other hand, both RNA-align and our RTM-align have good performance on scale consistency.

### RTM-align has superior performance on short RNA pairs

Our alignment program, RTM-align, selects the higher score between RTM-score with basic alignment (RTM-score_BA_) and RTM-score with fragment alignment (RTM-score_FA_) when the input pairs range from 20 to 50 nucleotides (Fig. 3a). The fragment alignment technique is specially introduced to improve the performance of short RNA pairs.

To evaluate the performance of our method on the short RNA pairs, we calculated the RTM-score for the 2,882 pairs within the curated hard-pairs dataset. This dataset includes pairs in which two RNA structures exhibit high sequence similarity (>80%) and low aligned RMSD (<2 Å) but receive extremely low TM-score values (<0.5) from RNA-align. As illustrated in Fig. 3b, our method, RTM-align, using RTM-score_BA_, produces higher scores on the hard-pairs dataset compared to RNA-align, indicating that our methods can identify more similar short RNA pairs than RNA-align. The overall distribution of scores is higher, with RNA-align’s 75th percentile falling below its classification threshold (0.39) derived from Rfam-50, whereas RTM-align shows the opposite trend. Furthermore, incorporating fragment alignment increases the distribution even more. This ablation study demonstrates that our two main improvements, post-standardization and fragment alignment, both enhance the performance of short RNA pairs.

The results in Table 1 provide a more quantitative assessment of this conclusion. We assume that all pairs in the hard-pairs dataset should be considered similar and evaluate a program’s accuracy on these difficult cases using its classification threshold on the Rfam dataset. We present results using both the Rfam-50 and Rfam-1000 thresholds; however, the former is more appropriate since all challenging cases have lengths ranging from 20 to 50 nucleotides. Under these conditions, it is evident that RTM-align using RTM-score_BA_ outperforms RNA-align, as indicated by a 43% improvement in accuracy. Consistent with the trend in Fig. 3b, incorporating fragment alignment provides an additional 7% improvement in accuracy.

**Table 1.** Performance of each method on the Rfam family assignment task and on the hard-pairs dataset.

| Method | Dataset | Threshold | Acc. (Rfam) | Acc. (Hard-pairs) |
| --- | --- | --- | --- | --- |
| RNA-align | Rfam-50 | 0.39 | 0.9284 | 0.4730 |
|  | Rfam-1000 | 0.35 | 0.9911 | 0.6785 |
| RTM-align w/o FA | Rfam-50 | 0.49 | 0.9362 | 0.9004 |
|  | Rfam-1000 | 0.47 | **0.9927** | 0.9160 |
| RTM-align | Rfam-50 | 0.50 | 0.9363 | 0.9707 |
|  | Rfam-1000 | 0.47 | **0.9927** | **0.9805** |

It is important to note that the fragment alignment procedure exclusively enhances RTM-align’s performance on extreme cases, such as those in the hard-pairs dataset, but does not yield higher scores for the majority of cases. To verify this, we compare the distribution of RTM-score_BA_ and RTM-score_FA_ on dataset Rfam-50 and hard-pairs. As shown in Fig. 3c, RTM-score_FA_ has a close distribution to RTM-score_BA_ on the hard-pairs dataset. In contrast, on Rfam-50 which is a dataset containing more general pairs, RTM-score_FA_ performs significantly worse. This difference in distribution indicates that applying fragment alignment is a stringent strategy that specifically targets extreme cases in the hard-pairs dataset, where global misalignment is caused by local structural mismatches of individual bases. In normal cases, such as those in Rfam-50, the scores are not influenced by using fragment alignment. The results indicate that fragment alignment is particularly effective in addressing local structural differences that lead to global misalignment in challenging cases.

### RTM-score has better correlation with other metrics in CASP15

The evaluation metrics for RNA structure prediction in the CASP15 competition, as demonstrated by Das et al. (31), show that the TM-score from RNA-align exhibits a notably lower correlation with other metrics for the short sequence R1117. Specifically, R1117 is categorized as an “easy” PreQ1 riboswitch target comprising only 30 nucleotides. Nevertheless, the TM-scores assigned by RNA-align to submissions for this sequence are unexpectedly low. For example, we visualized R1117 submission TS284_7, TS232_1, TS287_3, and TS270_3 (Fig. 4c), which are the top-1 submissions for R1117 in terms of RMSD, lDDT, GDT_TS and TM-score respectively. RNA-align’s TM-score for the first three submissions is only 0.371, 0.401, and 0.426 respectively. These scores are perilously close to the classification threshold of 0.35 derived from our Rfam-1000 dataset and are even lower than the threshold of 0.45 proposed in the foundational study by (21). Moreover, the top-1 submission suggested by RNA-align (TS270_3) scored notably worse than the other submissions in the other three metrics, for example, the RMSD of all the other three submissions is below 2.5 Å, while TS270_3 is above 4.0 Å. These discrepancies suggest that RNA-align’s method of assessing short RNA structure similarity might be biased, as it provides ambiguous judgments even when other metrics like RMSD, lDDT, and GDT_TS indicate a higher level of confidence in these cases.

**Fig. 4.**
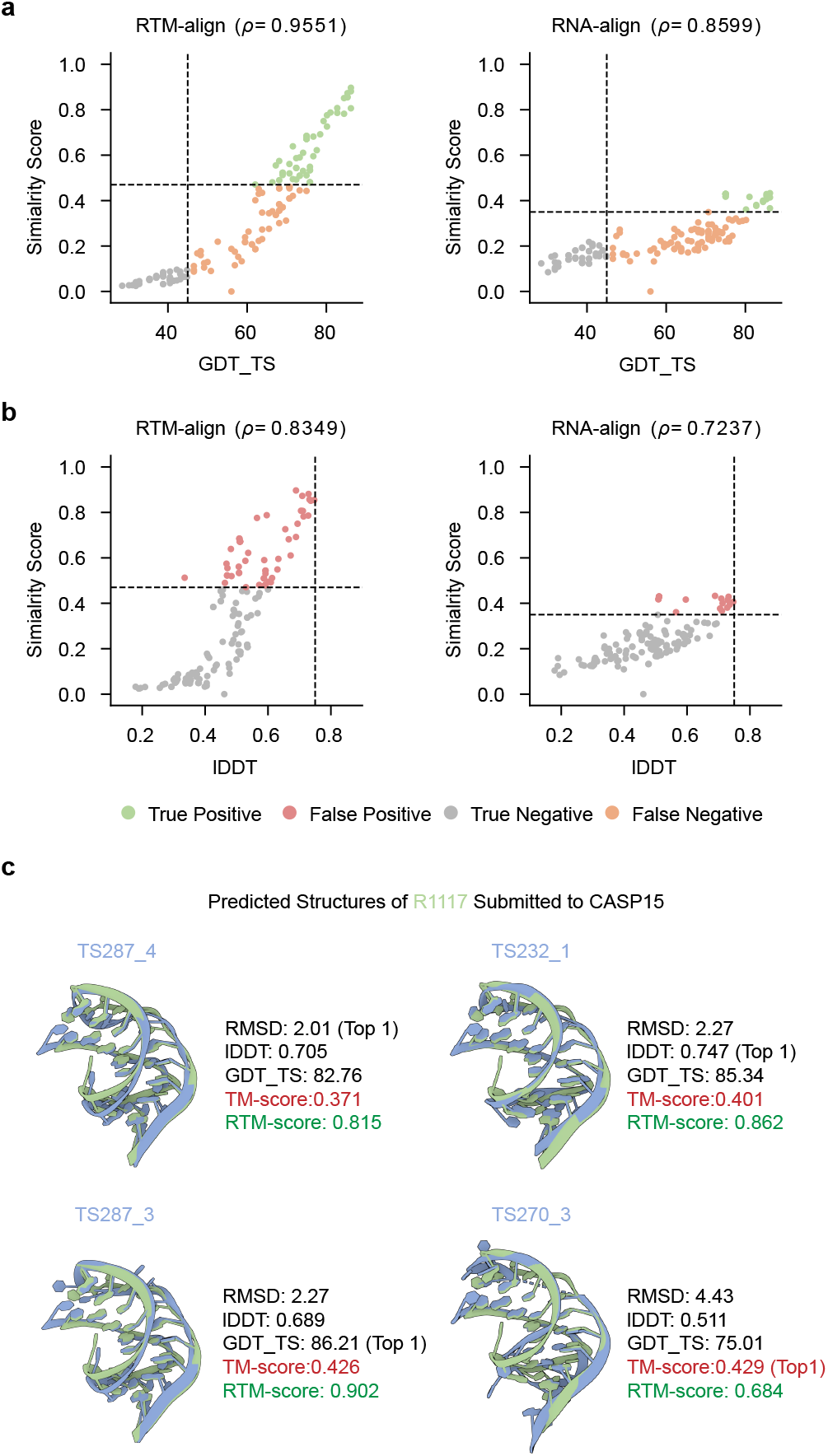
**(a)** Correlation of TM-score/RTM-score and GDT_TS **(b)** Correlation of TM-score/RTM-score and LDDT. **(c)** Performance comparison of RTM-score and TM-score on four submissions for R1117: TS284_7, TS232_1, TS287_3 and TS270_3

In contrast, RTM-score not only ensures uniform scoring scales but demonstrates a stronger correlation with other established scoring metrics. For the aforementioned cases, our program yields an RTM-score of 0.815 for TS287-4, 0.862 for TS232-1, 0.902 for TS287_3, and 0.684 for TS270_3, significantly surpassing the classification threshold of 0.47 for RTM-score, and TS270_3 is no longer the top-1 submission in terms of RTM-score. These scores align with the assessments provided by lDDT, GDT_TS, and RMSD for these submissions.

To evaluate the overall performance, we compared the score generated by RTM-align and RNA-align, as well as GDT_TS and lDDT, across all 153 submissions for case R1117 in CASP15. Spearman’s rank correlation coefficient was employed to assess the alignment between the two similarity metrics. As illustrated in 4(a)(b), RTM-score achieves a higher correlation score with GDT_TS and lDDT than RNA-align. Further, based on the similarity threshold of 0.45 for GDT_TS and 0.75 for lDDT provided by (31), we classified all submissions into positive and negative samples in conjunction with the thresholds set by RNA-align and RTM-score. We found that, on GDT_TS, RTM-score exhibits a higher true positive rate (TP) and a lower false negative rate (FN) (Fig. 4a). These trends were not mirrored with lDDT, which can be attributed to the extreme strictness of the lDDT threshold for R1117, as proposed in (31). These findings suggest that RTM-score provides more accurate and equitable scores, particularly for short RNAs, and stands as a promising candidate for a standard scoring metric in future CASP competitions.

## Discussion

In this paper, we presented RTM-align, an RNA structure alignment tool optimized for the structural comparison of short RNAs. Benchmarking results demonstrated that RTM-align significantly improved classification accuracy (from 68% to 98%) on the challenging short-pairs dataset compared to the well-known tool RNA-align. Furthermore, it maintained robust performance for longer RNAs, achieving consistent results regardless of RNA length. We also illustrated that RTM-align could provide a fair metric for evaluating RNA structure prediction methods, particularly for short RNA targets, such as R1117 in CASP15. Given the essential role of RNA structure comparison in understanding RNA functions and the critical importance of short RNAs in various biological pathways, RTM-align is poised to significantly enhance functional analysis in the field of RNA structural biology.

Although RTM-align offers promising advantages over existing RNA alignment tools, several limitations could be considered. The accuracy on the hard-pairs dataset did not reach 100% (Supplementary Fig. S7). The primary reason for errors in these cases could be the overly strict upper bound set for the penalty coefficient *β* in fragment alignment, which limited the score gains from fragment alignment. Additionally, these cases exhibited high RMSD both before and after fragment alignment, indicating widespread minor perturbations across most positions, a common challenge for all TM-score-based methods. While the current version of RTM-align provides accurate alignment evaluations for RNA, future versions will address these challenges to achieve more comprehensive structure comparisons.

## AUTHOR CONTRIBUTIONS

Z.Q., S.X. and T.S. conceived the project. Z.Q. and S.X. designed the methodology and conducted the experiments. S.S. supervised the study. Z.Q., S.X., and J.W. drafted the manuscript. All authors reviewed and approved the final version of the manuscript.

## CODE AVAILABILITY

The source code for RTM-align is available at https://github.com/BEAM-Labs/RTM-align.

## DATA AVAILABILITY

The pairs used for calculating the scaling factor can be accessed at https://doi.org/10.6084/m9.figshare.25903405.v1. Additional datasets, including Rfam-50, Rfam-1000, hard-pairs, and R1117, are also available at this URL.

## Supplementary Note 1: Fitting Details of Correlation Factor and Radius

To remain consistent with previous work(24), we employed a second-order exponential polynomial to fit the correlation factor, which follows the general form 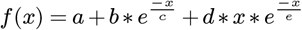. The final function is 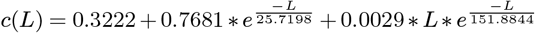.

Our fitting results for the radius indicate that it is proportional to *L*^0.3410^, whereas previous research on protein data suggests that this metric should be proportional to *L*^0.39^(24). RMalign(20) also reports a value of 0.39 for RNA data. However, due to the limited scope of our literature review, no one else has accurately measured this parameter for RNA, making it difficult to determine if 0.3410 is a biased result. Considering that we used more data points than RMalign for our fitting, our results should theoretically be more accurate, although inherent biases in the dataset cannot be completely eliminated. Nevertheless, any potential errors in our fitting of the radius will be minimized during the post-standardization process, so this fitting step should not pose significant issues.

**Fig. S1.**
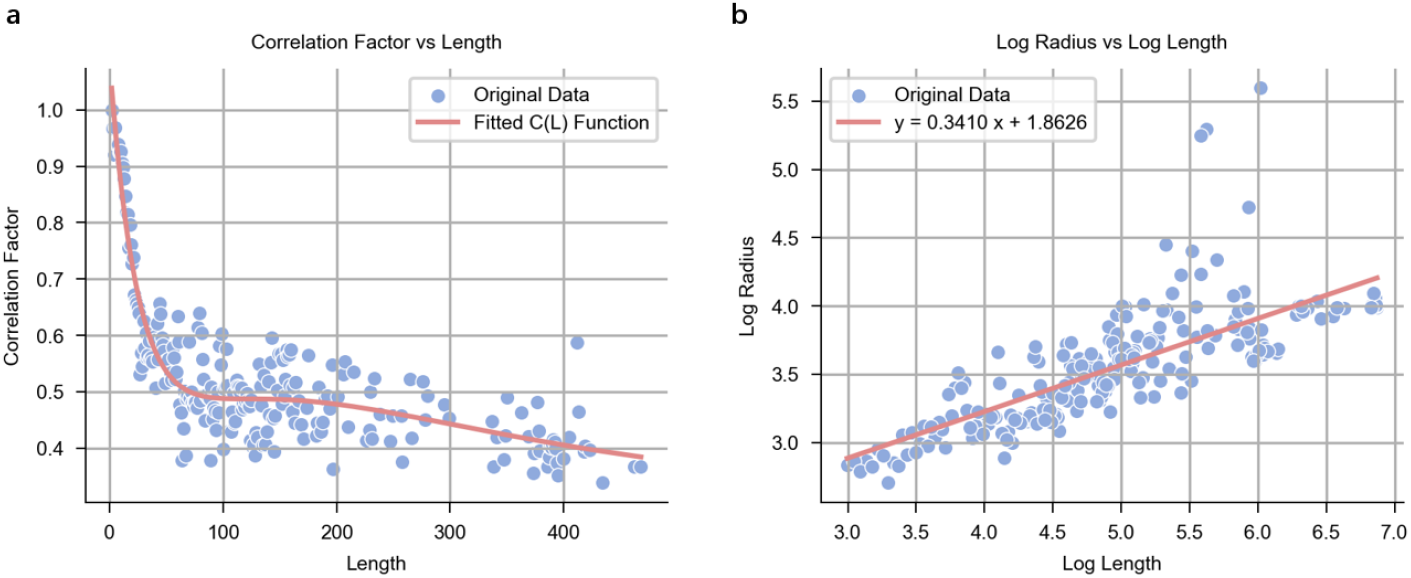
**(a)** Average correlation factor at each length and the fitted curve *c*(*L*), calculated on the random pairs dataset. **(b)** Average radius at each length and the fitted curve *R*(*L*), calculated with the unique RNA structures in the random pairs dataset.

## Supplementary Note 2: Fitting Details of mean(L) and std(L)

**Fig. S2.**
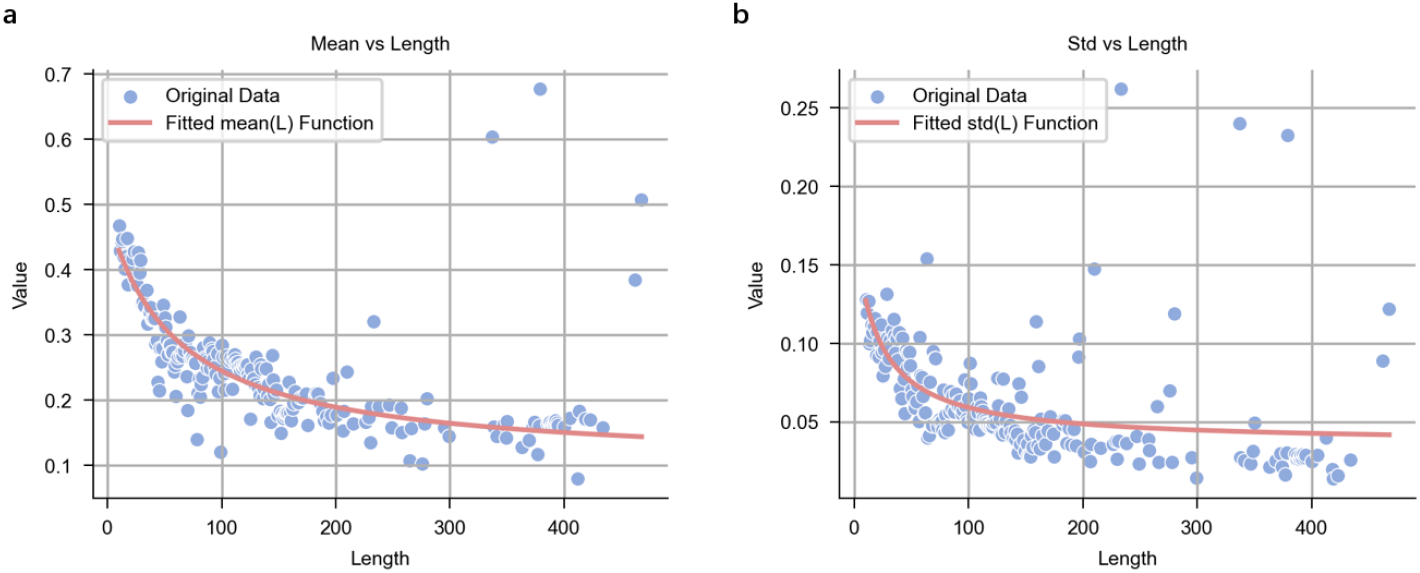
**(a)** Mean of TM-score calculated with the derived *d*_0_(*L*) for RNA at each length, and the fitted curve *mean*(*L*), calculated on the Random Pairs dataset. **(b)** Standard deviation of TM-score calculated with the derived *d*_0_(*L*) for RNA at each length, and the fitted curve *std* (*L*), calculated on the Random Pairs dataset.

## Supplementary Note 3: Finding the Upper Bound of the Punishment Coefficient

**Fig. S3.**
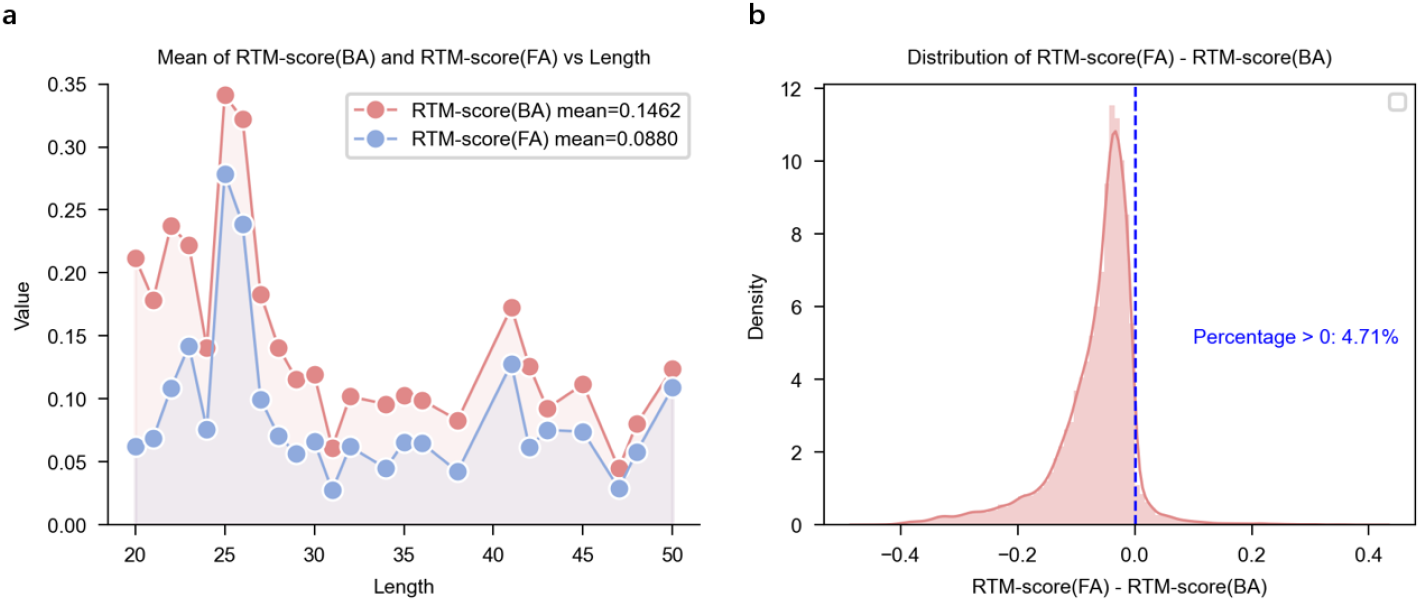
**(a)** Comparison of the scale of RTM-score_BA_ and RTM-score_FA_ in the length interval of [20, 50], calculated on the Rfam-50 dataset. **(b)** Advantage of RTM-score_FA_ over RTM-score_BA_ on the Rfam-50 dataset.

## Supplementary Note 4: The Functional of Positional Bias

**Fig. S4.**
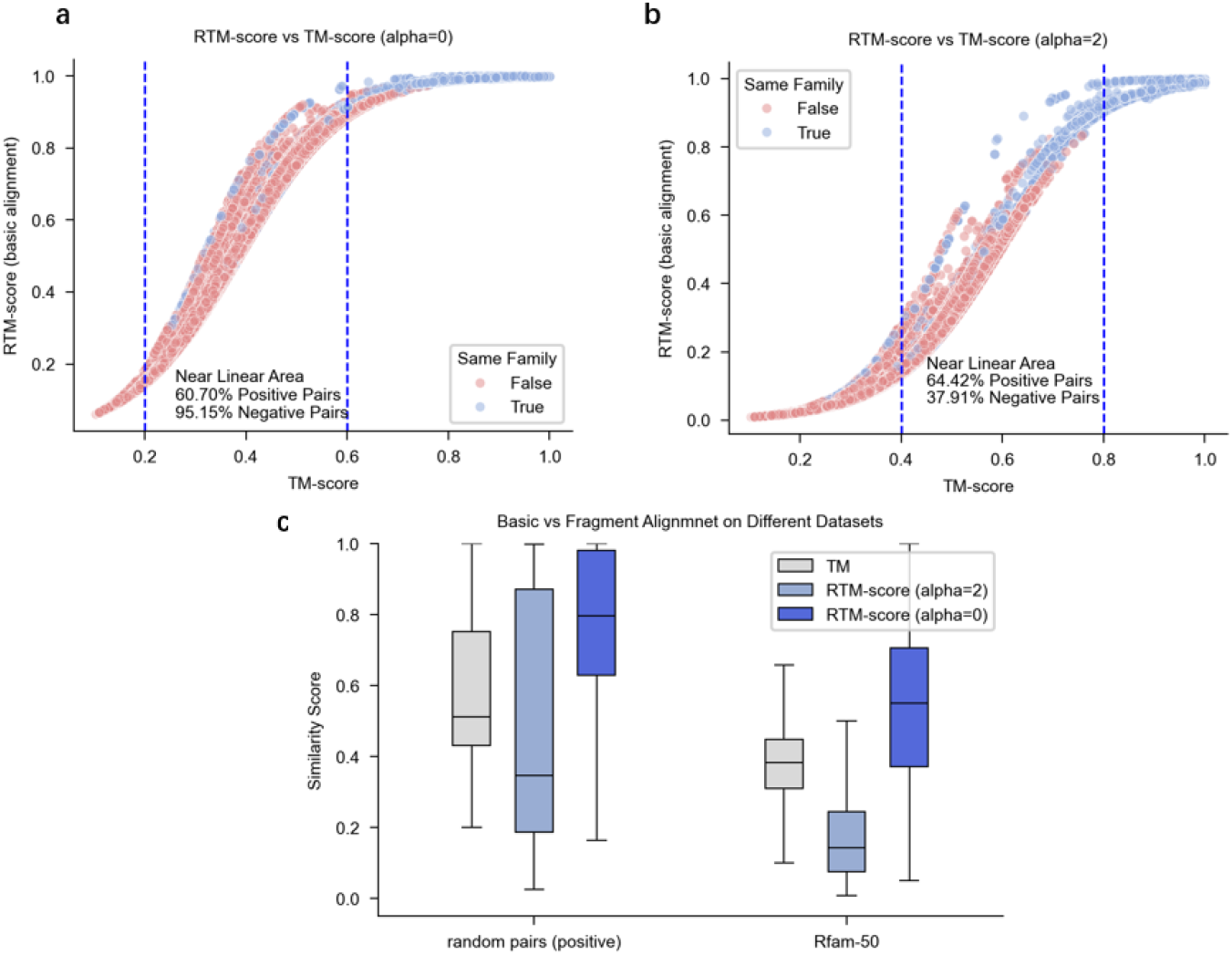
**(a)** Each pair’s RTM-score_BA_ and TM-score in Rfam-50 dataset, when *α* = −2. **(b)** Each pair’s RTM-score_BA_ and TM-score in Rfam-50 dataset, when *α* = 0. Red pairs stand for negative pairs and blue pairs stand for positive pairs. **(c)** Distribution of TM-score, RTM-score_BA_ (*α*=-2) and RTM-score_BA_ (*α*=0) on Random Pairs dataset and the positive pairs of Rfam-50 datasets.

## Supplementary Note 5: Slope of Scale

**Table S1.** Slope of average score vs length at intervals of [20, 50], [20, 80], [20, 100], and [20, 200].

| Method | 20-50($\times 10^{-3}$) | 20-80($\times 10^{-3}$) | 20-100($\times 10^{-3}$) | 20-200($\times 10^{-3}$) |
| --- | --- | --- | --- | --- |
| RNA-align | 0.4525 | 0.1703 | 0.3858 | 0.1611 |
| RMalign | 5.9630 | 3.2425 | 2.1071 | 0.9116 |
| RTM-align | 2.9380 | 1.6054 | 0.4990 | 0.0625 |

## Supplementary Note 6: Discussion of the Causes of Hard Pairs

The prime intuition of our work is to fix RNA-align’s bias on short RNAs, and the thing that helps us notice RNA-align’s shortcoming is the discovery of ’hard pairs’ while using the program. The so-called ’hard pairs’ are pairs of RNAs that have high sequence similarity (>0.9), and some of them are even different pdb files for the same sequence, but these pairs are given extremely low structure similarity scores (<0.4) by RNA-align, which is counterintuitive. We note that this phenomenon only happens on short RNA pairs ranging in [20, 50]. Additionally, the aligning procedures should not be the problem, because most residuals of most hard pairs are aligned.

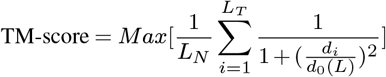

TM-score is calculated as the formula above, standing from it we raise two possible explanations for RNA-align’s biased performance on hard pairs. The first explanation is: the fitted *d*_0_(*L*) of RNA-align is unreasonably small on short RNAs, thus not only causes the result TM-score to be small, but also makes RNA-align over-sensitive to substle changes in the distances between aligned residuals *d*_*i*_. From Supplementary Fig. S5 we can see that RNA-align’s *d*_0_ has the smallest value among RNA-align, RMalign and RTM-score, when the length is fixed between 20 to 50. This also explains why RMalign gives higher scores on this length interval.

**Fig. S5.**
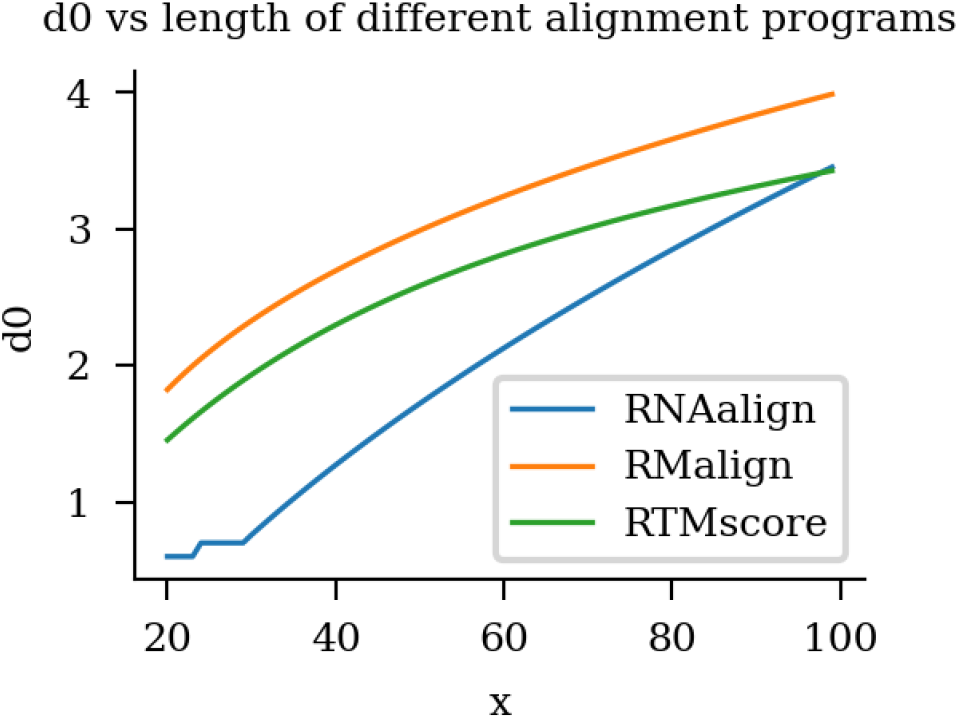
*d*_0_ vs length of RNA-align, RMalign and RTM-score respectively.

Another possible explanation mainly considers the aligning procedure. We believe some of the hard pairs are misaligned globally mainly because of some local deviations. For example, one tiny deviation at a single residual may cause the whole structure to have a global shift. This shouldn’t be a matter to pairs of ordinary lengths cause the overall shift is tiny. But when dealing with short RNAs, these shifts can’t be overlooked anymore since the *d*_0_ at these lengths are originally small and any small change in *d*_*i*_ may cause non-neglectable change in similarity score. To verify this, we compare the RMSD of the aligned part of structure pairs before and after fragment alignment (Supplementary Table.S3). Pairs in Rfam-50 have an average RMSD decrease of 35.71% while that of hard-pairs dataset is 42.11%. This means that pairs in hard-pairs can be intrinsically aligned closer than ordinary pairs when neglecting several local misalignments.

## Supplementary Note 7: More Visualizations of Short RNA Pairs with RTM-score

**Fig. S6.**
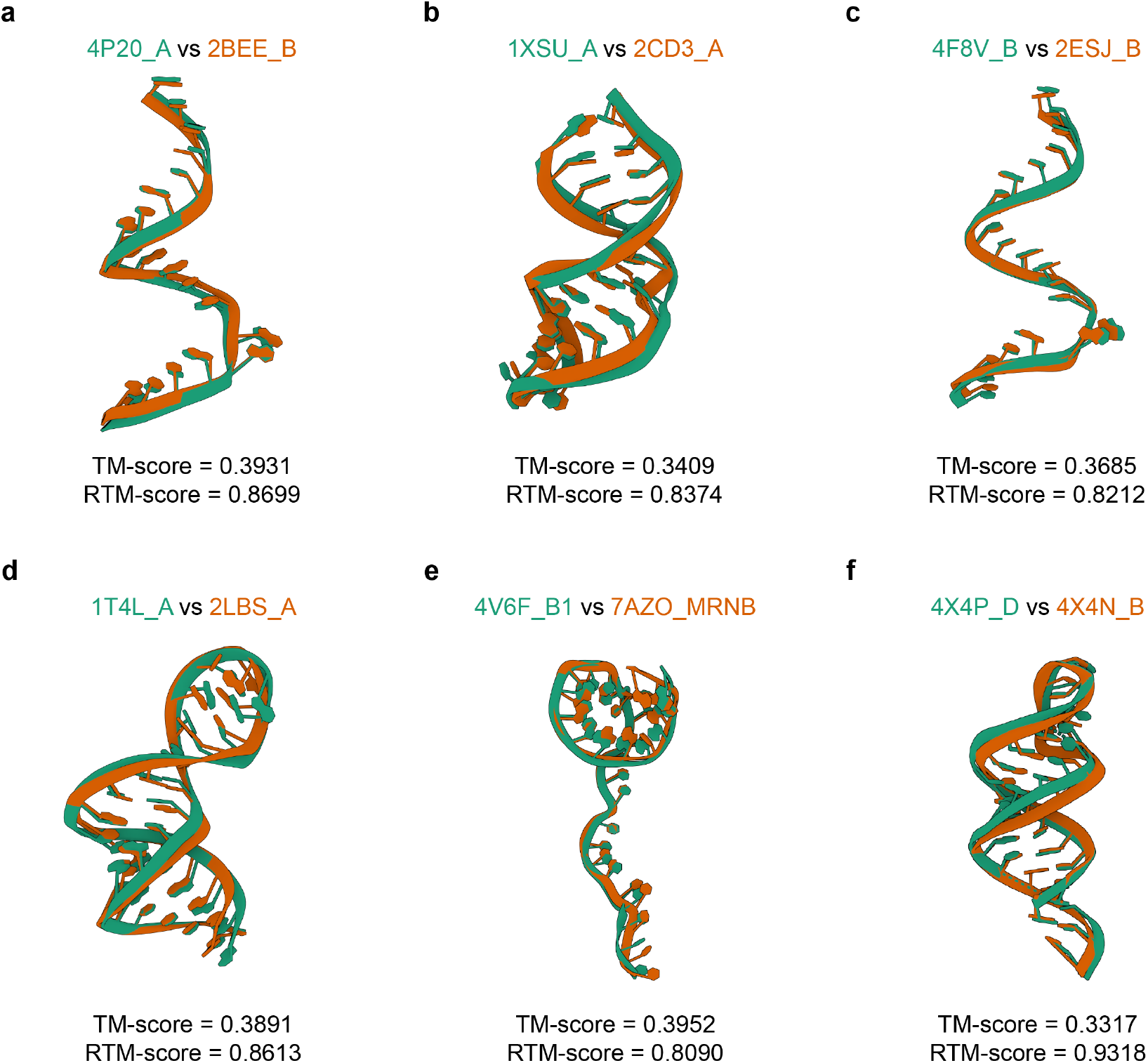
More visualizations of short RNA pairs with RTM-score.

## Supplementary Note 8: Limitations of RTM-align on Some Challenging Pairs

Although RTM-align offers promising advantages over existing RNA alignment tools, several limitations must be considered. Our accuracy on the hard-pairs dataset did not reach 100% (Supplementary Fig. S7). The primary reason for errors in these cases is the overly strict upper bound for the penalty coefficient *β* in fragment alignment. This strict upper bound limited the score gains from fragment alignment. Our intention was to minimize the impact of fragment alignment on cases outside the bad-case set. However, noise exists in the dataset; for instance, we cannot ensure that Rfam-50 does not contain pairs similar to the hard-pairs. Therefore, determining a more reasonable penalty size remains an area for improvement.

Additionally, the cases where we failed share a common characteristic: the RMSD remains high after fragment alignment compared to other pairs in the hard-pairs dataset. These three cases show less than a 25% decline in RMSD after fragment alignment (Supplementary Table.S2), while the average decline for the hard-pairs dataset is above 40% (Supplementary Table.S3). This suggests that their structural misalignment is not due to a few misplaced bases but rather widespread minor perturbations across most positions. The current fragment alignment approach cannot effectively address this issue. This type of case presents a significant challenge for all methods based on TM-score.

**Fig. S7.**
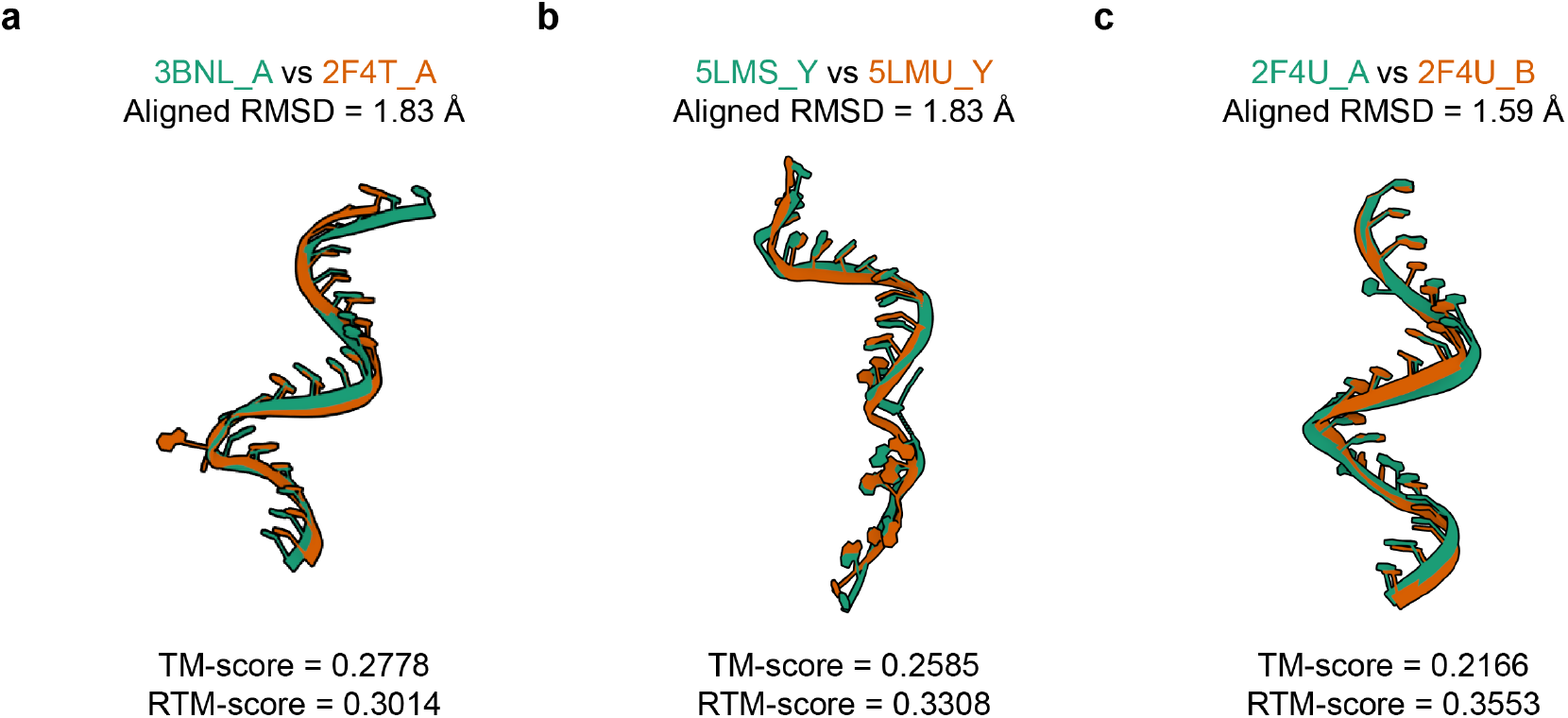
In some similar RNA structure pairs, RTM-align failed to have satisfying performance.

**Table S2.** The RMSD of the aligned part of pair 2F4U_A and 2F4U_B, 3BNL_A and 2F4T_A, 5LMS_Y and 5LMU_Y before and after fragment alignment. Delta here is the percentage of declined RMSD value after fragment alignment.

| RNA Pair | $L_{\min}$ | RMSD | RMSD<sub>frag</sub> | Delta (%) |
| --- | --- | --- | --- | --- |
| 2F4U_A 2F4U_B | 21 | 1.59 | 1.20 | 24.53 |
| 3BNL_A 2F4T_A | 21 | 1.83 | 1.58 | 13.66 |
| 5LMS_Y 5LMU_Y | 20 | 1.83 | 1.59 | 13.11 |

**Table S3.** The average RMSD of the aligned part of structure pairs before and after fragment alignment. Delta here is the percentage of declined RMSD value after fragment alignment, calculated on Rfam-50 and hard-pairs datasets. Columns with L=20,21 mark the statistics on RNA pairs with lengths of 20 or 21.

| Dataset | RMSD | RMSD<sub>frag</sub> | Delta (%) | RMSD (L={20,21}) | RMSD<sub>frag</sub> (L={20,21}) | Delta (%) (L={20,21}) |
| --- | --- | --- | --- | --- | --- | --- |
| Rfam-50 | 2.38 | 1.53 | 35.71 | 1.92 | 1.43 | 25.52 |
| Bad-case | 1.33 | 0.77 | 42.11 | 1.25 | 0.72 | 42.40 |

